# Dietary protein source dictates the impact of obesogenic diets on hepatic steatosis and insulin resistance via carnitine-dependent regulation of acetyl-CoA carboxylase

**DOI:** 10.64898/2026.06.25.732886

**Authors:** Frédéric Bégin, William Gagnon, Lia Rossi Perazza, Patricia L Mitchell, Thomas Guerbette, Bertrand Bouchard, Michaël Shum, Alexandre Caron, Christine Des Rosiers, Stanislaw Deja, Phillip J. White, André Marette

## Abstract

Nutritional strategies to mitigate obesity and type 2 diabetes (T2D) have largely focused on dietary fat and carbohydrate composition, with less attention given to protein sources. While total dietary protein intake is recognized as an important modulator of energy balance and glucose metabolism, it remains unclear how the composition of dietary proteins can influence energy metabolism and body weight gain. Here, we investigated the metabolic effects of three distinct protein sources from meat (pork), dairy (casein) and plant (soy) on either a low-fat low sucrose (LFLS) or a high-fat high sucrose (HFHS) diet. While protein sources failed to influence metabolic homeostasis on LFLS, mice kept on the HFHS diet were distinctly impacted by the dietary protein sources. Pork and to a lesser extent soy protein feeding exacerbated obesity, glucose intolerance, and hepatic insulin resistance. Remarkably, livers of mice fed pork or soy protein on the HFHS diet were characterized by extensive microvesicular steatosis compared to the predominant macrovesicular steatosis in HFHS fed mice fed casein protein. Liver transcriptomic and metabolomic signatures in pork and soy protein fed mice were consistent with increased mitochondrial beta-oxidation. Intake of pork and soy proteins in HFHS fed mice lead to a striking reduction in hepatic acetyl CoA carboxylase 2 (ACC2) protein levels relative to casein fed HFHS mice. Pork and soy feeding raised carnitine exposure in the post-prandial period and we determined that exposure of hepatocytes to carnitine provokes downregulation of ACC2 and hepatic insulin resistance in the presence of palmitate:oleate and fructose. Collectively, these findings identify a novel mechanism by which dietary proteins modulate obesity and associated metabolic disturbances through a carnitine-mediated regulation of ACC2 protein and mitochondrial lipid handling in liver.

**GRAPHICAL ABSTRACT:** 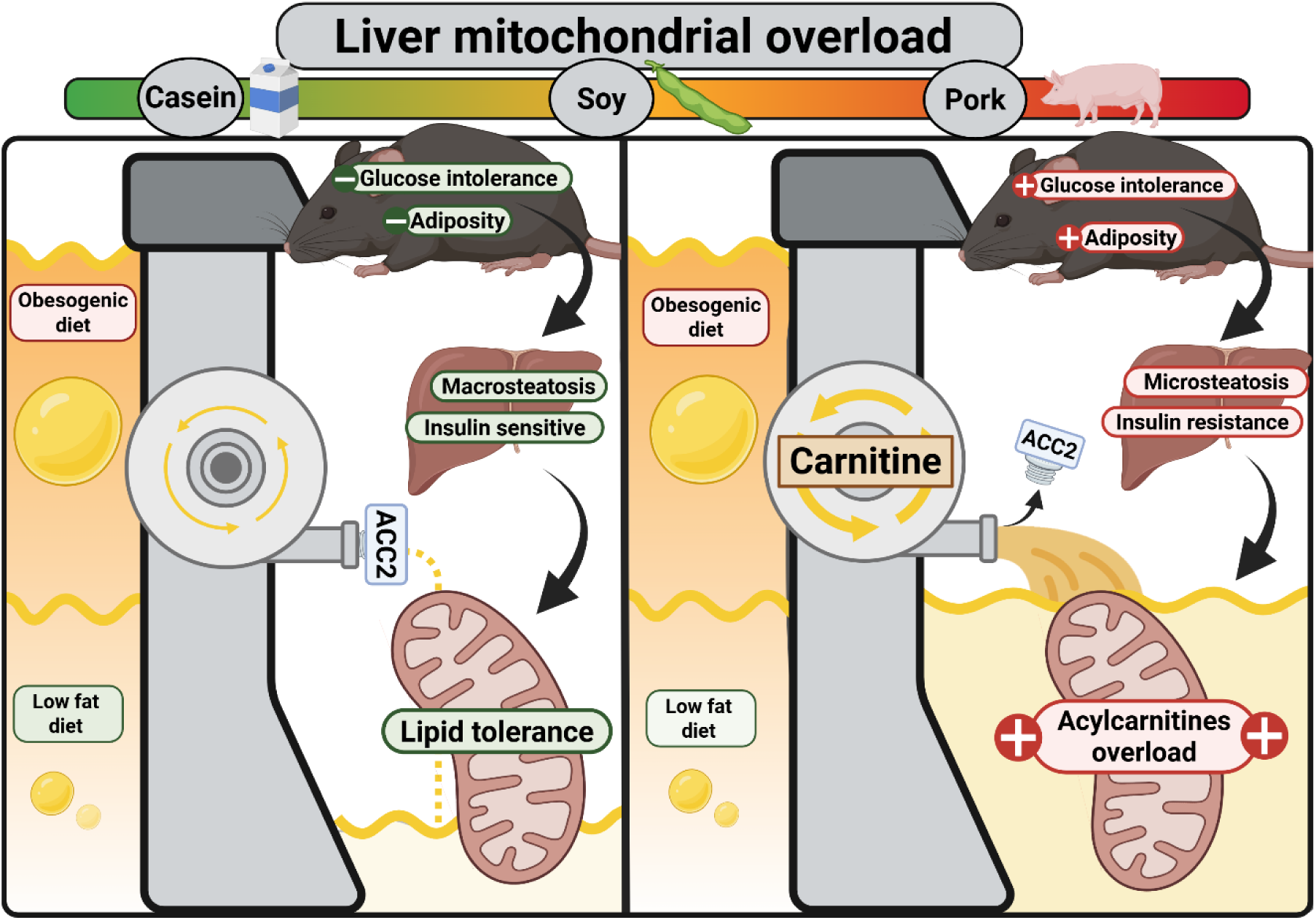

## INTRODUCTION

Obesity and type 2 diabetes (T2D), characterized by excess adiposity, hyperglycemia, dyslipidemia and insulin resistance, are major public health concerns worldwide^1–3^. Nutritional strategies aimed at targeting these conditions have historically focused on dietary lipids and carbohydrates, which are widely viewed as the primary nutritional drivers of metabolic diseases. In contrast, the role of dietary protein as a potential contributor to the development of obesity and T2D has historically been largely overlooked.

Most previous studies have examined total protein intake (high versus low protein diets), with limited consideration of protein source-specific effects^4–11^. Although increasing total protein can improve metabolic outcomes in some contexts, long-term adherence to high protein diets remains highly variable, and compliance is a key determinant of their effectiveness^12,13^. Modifying protein sources, rather than total protein quantity, may therefore represent a more feasible and sustainable nutritional strategy.

Emerging evidence from preclinical studies indicates that substituting casein with alternative, more human-relevant, protein sources can markedly influence susceptibility to diet-induced obesity^14–18^. However, there is still no clear mechanistic framework explaining how different protein sources influence chronic disease risk through their interaction with obesogenic diet. This is critically important since animal studies indicate that simply changing protein source alone can produce metabolic effects comparable in magnitude to those achieved by altering total protein content^16^.

Recently, we reported that a mixed protein diet designed to reflect the composition of the human Western diet exacerbates high-fat high sucrose (HFHS)-induced obesity and related metabolic disturbances compared to a casein-based diet, still often used in nutritional studies with murine models^19^. Importantly, these metabolic effects were only observed in mice fed a HFHS diet, but not in those fed a low-fat low sucrose (LFLS) diet, indicating that dietary protein composition primarily influences metabolic homeostasis in the context of obesogenic diet intake. Feeding diverse protein sources was found to rapidly reshape the gut microbiota and to promote postprandial accumulation of hepatic acylcarnitines (AC), a hallmark of metabolic diseases reflecting a bottleneck in mitochondrial fatty acid oxidation that is associated with insulin resistance^20^. While some of the metabolic effects of the mixed protein diet were gut microbiota dependent, we observed that the increase in circulating and hepatic AC persisted even after antibiotic-mediated microbiota depletion^19^. Therefore, the molecular mechanism by which dietary protein impact liver lipid metabolism remains unresolved.

Here we report that dietary pork, and to a lesser extent soy protein, exacerbates HFHS-induced weight gain, glucose intolerance, and hepatic insulin resistance compared with casein. Our data support a conceptual model in which dietary protein source modulates susceptibility to HFHS-induced metabolic dysfunction by altering insulin signaling and mitochondrial lipid oxidation in liver. Remarkably, pork protein exerts this effect through a carnitine-dependent down-regulation of ACC2 abundance. This model provides a mechanistic explanation to the previously unresolved question of how dietary protein sources interact with obesogenic diet to promote hepatic AC accumulation and liver insulin resistance, identifying carnitine as a readily modifiable factor regulating hepatic metabolism in obesity and T2D.

## RESULTS

### Dietary pork and soy protein exacerbate obesity, glucose intolerance and liver insulin resistance in HFHS-fed mice compared to casein

To determine whether dietary protein sources modulate obesity and metabolic homeostasis, we selected casein (C), soy (S) and pork (P) proteins as representative of dairy-, plant- and meat-derived protein sources, respectively. Each protein was incorporated at 15% of total caloric content into either a HFHS (H) or LFLS (L) diet, generating 6 experimental diets (HP, HS, HC, LP, LS, LC). All diets were isocaloric and matched for macronutrient composition (Figure 1A), fiber content, and saturated-to-polyunsaturated fat ratio (Supplementary table 1 to 3).

**Figure 1:**
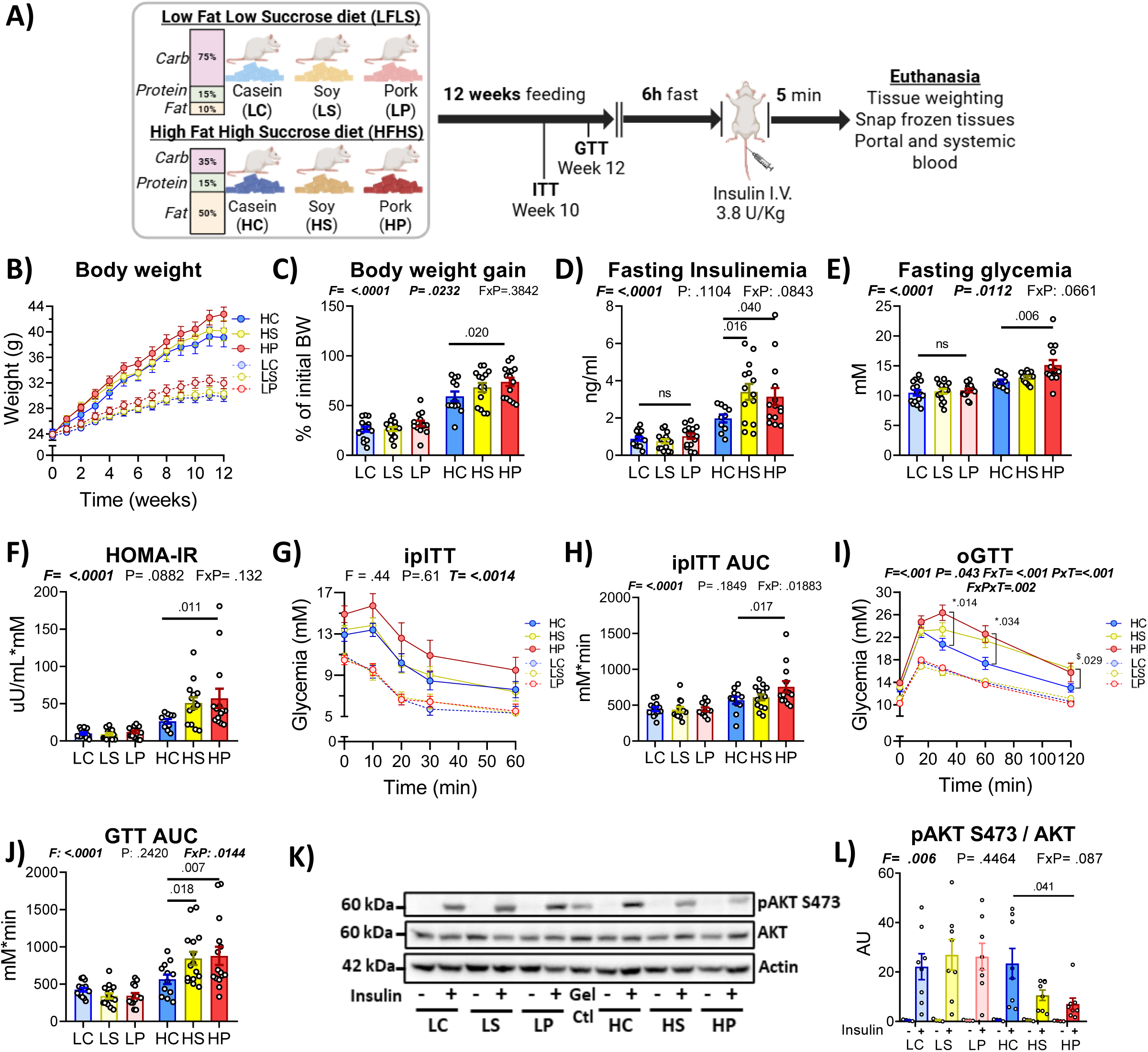
Dietary pork and soy protein exacerbate obesity, glucose intolerance and liver insulin resistance in HFHS-fed mice compared to casein. **A-C**) Mice (n=12-15) were fed for 12 weeks with diet containing 15 % of the total caloric content as protein in either high fat high sucrose (HFHS) diet or low-fat low sucrose (LFLS) diet. Protein sources were either isolated casein (HC and LC), isolated soy (HS and LS) or lyophilised pork (HP and LP). Body weight was measured weekly (B) and body weight gain (C) was calculated comparing the final weight after 12 weeks to the weight at baseline. **D-F**) Fasting insulinemia (D) was measured (n=10-15) by ELISA using 6h fast tail vein plasma collected at time 0 during insulin tolerance test and were used with fasting glycemia (E) to derive HOMA-IR index (F) corrected for mice (fasting insulin(uU/mL)*fasting glucose(mM)*/22,5). **G-H**) Intraperitoneal insulin tolerance test (ipITT) was determined on the 10^th^ week (n=11-14) and area under the curve (AUC) was calculated from baseline value for each animal (H). **I-J**) Oral glucose tolerance test (oGTT) was performed during the 12^th^ week (n=12-15) and area under the curve (AUC) was calculated from baseline value for each animal(J). Stars (*) refer to the result of Dunnet’s post-hoc for HC to HP comparison, and dollars ($) sign refer to HC to HS comparison **K**) Phosphorylated AKT on serine 473 in liver (Insulin n=8, Saline n=4), corrected for total AKT, and using actin as for loading control. Gel Ctl= Gel loading control. **L**) Quantification of the blots in J. Intensity of the bands were normalised on actin and gel control to give relative intensity. Data are represented as means ± s.e.m. Statistical analyses were performed using either two-way ANOVA, a three-way ANOVA using a mixed model effect for repeated measures in time (ipITT and oGTT), followed by a Dunnet’s post-hoc test for comparison to casein group. P-values of general effect for diet fat and sucrose (F), protein (P), time (T) are recorded under the title of each graph, followed by the p-values of the corresponding factor interaction effects.

Under LFLS conditions, protein sources had no impact on body weight or fat mass gain (Figure 1B and Supplementary figure 1A). In contrast, under HFHS conditions, mice fed pork protein showed an increase in body weight gain compared to casein-fed controls (Figure 1B and C), accompanied by increased fat mass, mostly explained by epididymal white adipose tissue (WAT) mass (Supplementary Figure 1A and B). HFHS mice fed soy protein showed an intermediate phenotype. There was no significant effect of the dietary protein source on other WAT depots, gastrocnemius or liver weight but a small but significant effect of soy protein intake on brown adipose tissue (BAT) of HFHS-fed mice was observed (Supplementary Figure 1C-H).

Similarly, glucose homeostasis was unaffected by the protein source under LFLS conditions (Figure 1E-J). However, after 10 weeks of HFHS feeding, clear interactions between the obesogenic diet and protein source emerged. Mice fed pork protein exhibited significantly elevated fasting glycemia and insulinemia compared to casein-fed HFHS mice (Figure 1D and E), resulting in a significant increase in HOMA-IR (Figure 1F). Soy protein did not alter fasting glucose levels but increased fasting insulin relative to casein-fed HFHS mice. Insulin sensitivity, as assessed by an *i.p.* insulin tolerance test, tended to be impaired in HFHS-fed mice consuming pork protein (Figure 1G and H), however oral glucose tolerance tests (oGTT) revealed markedly impaired glucose tolerance for this group, and to a lesser extent for soy protein-fed mice, as shown by increased glucose excursion (Figure 1I and J). Glucose-stimulated insulin secretion (GSIS) during the oGTT was similarly elevated across all HFHS groups compared to LFLS controls, with no effect of protein source (Supplementary Figure 1I and J). Insulin-stimulated Akt phosphorylation in the liver was reduced in HFHS-fed mice fed pork protein, and to a lesser extent soy protein, compared to casein-fed mice (Figure 1J and K). Taken together, these results demonstrate that dietary protein sources dictate the detrimental impact of an obesogenic diet, with pork-derived protein exacerbating the impact on weight gain, glucose tolerance and hepatic insulin resistance more than soy.

### Dietary pork and soy protein promote microvesicular hepatic steatosis and increased lipid mobilization in HFHS-fed mice

Given the more severe hepatic insulin resistance observed in HFHS-fed mice consuming pork or soy protein compared to casein, we next examined if protein source influenced systemic and liver lipid metabolism after 12 weeks of dietary treatment. Liver and plasma triglyceride and cholesterol content were unaffected by dietary protein sources under HFHS feeding (Figure 2A and Supplementary 2A-B). However, under LFLS conditions, pork protein intake increased both hepatic and circulating cholesterol levels (Figure 2B and supplementary 2B). There was no effect of dietary protein source on serum content of the liver transaminases, AST or ALT, as well as inflammatory and fibrosis score in HFHS feeding (Supplementary Figure 2C-H). With the notable exceptions of interleukin-1b and Keratinocyte Chemoattractant (KC), most of the fibrosis and inflammatory markers were not affected by protein sources in HFHS fed animals (Supplementary Figure 2I).

**Figure 2:**
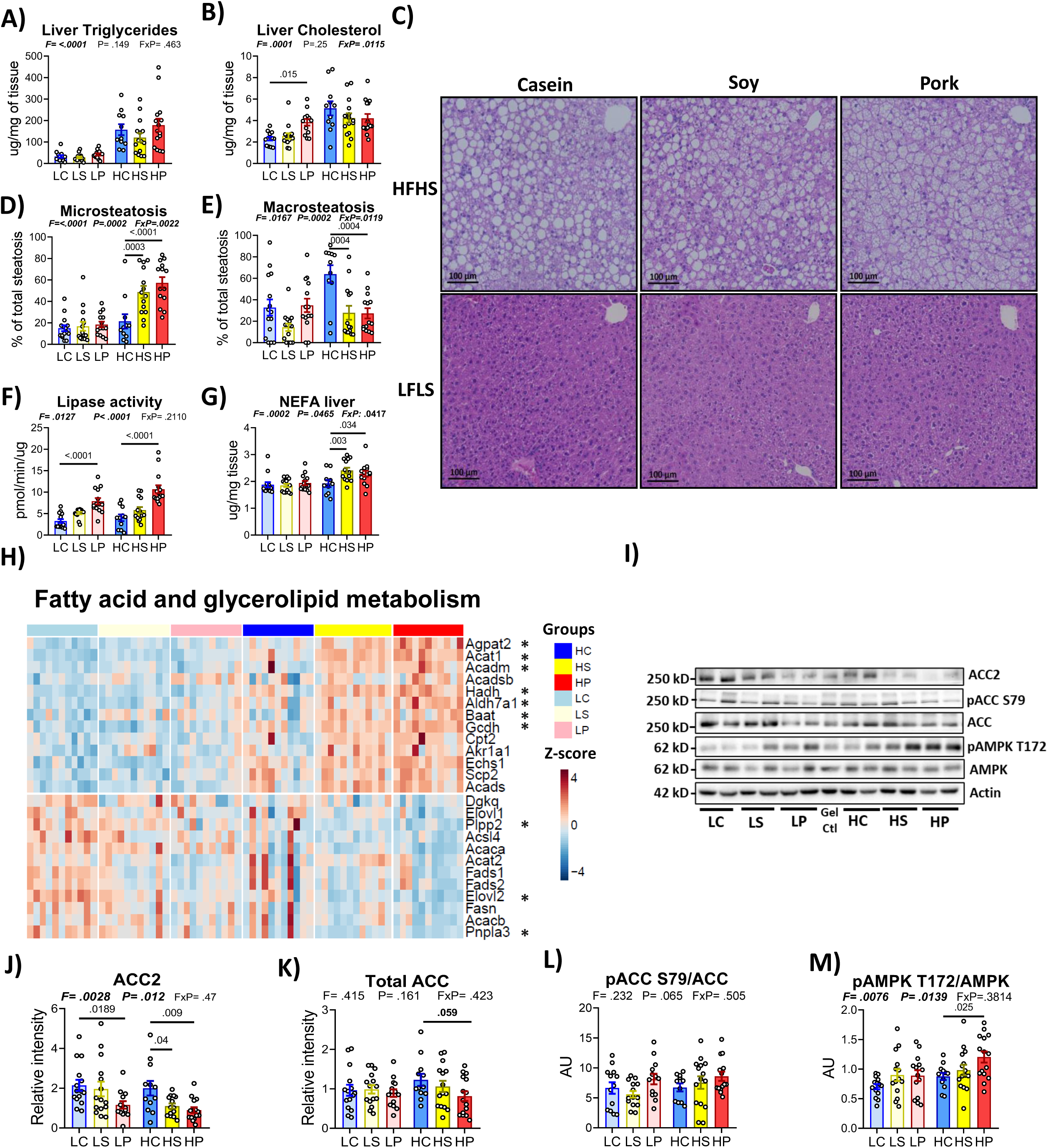
Chronic pork and soy consumption results in liver microsteatosis, transcriptional rewiring towards beta-oxidation and modulation of AMPK-ACC axis. **A-B)** Liver total triglycerides(A) and cholesterol (B) normalised to g of tissue weighted. **C**) Representative picture of HE staining slides of liver from mice (n=11-15) fed for 12 week and sacrificed after 6h fast. **D-E**) Mean of blinded visual quantification of three observers for micro (D) and macro (E) steatosis. **F)** Lipase activity measured by fluorescence plate kinetic assay of liver lysate. **G**) Total non esterified fatty acid in liver lysate **H)** Trimmed Mean of M-values (TMM, normalised feature counts) were used to visualise gene expression for significant differently expressed gene(padj<0.1) found during DESeq analysis for Pork to Casein comparison (overall protein effect, adjusted for diet background). Selection was made for gene with tag “Fatty acid” or “Glycerolipid” in their KEGG annotation, and stars (*) indicates that these genes were significant for the subset HFHS-Pork to HFHS-casein comparison. **I)** Representative blot for the indicated protein and their phosphorylation site, with 2 mice per group. Gel Ctl= Gel loading control. **J**-**M**) Quantification of the blots in K. Intensity of the bands were normalised on actin and gel control to give relative intensity. AU stands for Arbitrary units (ratio). Data are represented as means ± s.e.m. Statistical analyses were performed using a two-way ANOVA, followed by a Dunnet’s post-hoc test for comparison to casein group. P-values of general effect for diet fat and sucrose (F), protein (P) are recorded under the title of each graph, followed by the p-values of the corresponding factor interaction effects.

Despite no effect of dietary protein source on total hepatic lipid content in HFHS-fed mice, histological analysis revealed striking differences in the steatotic phenotype. Hepatocytes from HFHS-fed mice consuming pork or soy protein contained numerous small lipid vesicles, consistent with microvesicular steatosis, whereas hepatocytes from casein-fed mice predominantly contained larger lipid droplets that are characteristic of macrovesicular steatosis. (Figure 2 C-E). This shift in lipid droplet architecture suggests that dietary protein source influences intracellular lipid handling rather than total lipid storage. Accordingly, total hepatic lipase activity increased in the liver of pork protein-fed mice, suggesting higher triglycerides hydrolysis to non-esterified free fatty acids (NEFA) (Figure 2F). This effect was specific to pork protein regardless of background diet. However, hepatic NEFA accumulation was only observed under HFHS conditions, and occurred in both pork- and soy-fed mice (Figure 2G). Importantly, the similar magnitude of NEFA accumulation and microvesicular steatosis observed in both pork- and soy-fed HFHS mice suggests that increased lipase activity alone does not fully account for this phenotype. Instead, accumulation of NEFA in hepatocytes suggests a more downstream bottleneck in free fatty acid utilization.

### Consumption of pork and soy protein induces hepatic transcriptional rewiring toward lipid oxidation, mitochondrial respiration and oxidative stress buffering

We hypothesized that the major switch in lipid droplet morphology observed in HFHS-fed mice consuming pork or soy protein reflects altered intracellular lipid trafficking and/or changes in mitochondrial fatty acid oxidation. This prompted us to conduct liver transcriptomic analysis (RNAseq) to more precisely identify the molecular pathways involved. Differential expressed sequence (DESeq) analysis was performed using a factorial design incorporating both protein source (Casein, Soy and Pork) and diet (HFHS vs LFLS), accounting for dietary background when estimating protein-specific effects. To specifically interrogate pathways related to lipid handling, from lipid droplet dynamics to mitochondrial beta-oxidation, we focused on differentially expressed genes annotated with “fatty acid” or “glycolipid” terms in the KEGG database.

Genes that were significantly differently expressed between pork- and casein-fed HFHS groups were related to increased mitochondrial lipid oxidation (increased expression of *Acsl1*, *Hadh*, *Acadm* and *Acat1*), phosphopholipid remodeling (decreased *Plpp2* and increased *Agpat2*), reduced polyunsaturated fat synthesis (decreased *Elovl2*) and increased bile acid-glycine conjugation (increased *Baat*) (Figure 2H). *Pnpla3,* whose human ortholog carries the rs738409 single nucleotide polymorphism, is among the most strongly associated genetic contributors to MASLD development^21–23^. This gene was the most downregulated by pork protein feeding. Its downregulation is consistent with the microvesicular steatosis phenotype observed in these animals, as *Pnpla3* protein prevents triglyceride lipase access to the lipid droplet, therefore impacting its fragmentation^22^. Several genes were similarly regulated by soy protein, when compared to casein-fed HFHS animals, including *Pnpla3*, *Elovl2*, *Acat1*, and *Baat* (Supplementary figure 3A). Interestingly, pork protein feeding also led to upregulation of an array of genes involved in the detoxification or buffering of oxidative stress (Supplementary Figure 3B), an effect consistent with higher levels of reduced glutathione in liver of pork protein fed mice compared to soy and casein fed mice in HFHS diet (Supplementary figure 3C). Finally, pork protein feeding led to increased expression of multiple components of the mitochondrial respiratory chain. This is consistent with enhanced mitochondrial respiration, which is predicted to increase during higher lipid oxidation. Thus, both soy and pork protein ultimately lead to increased genes involved in the regulation of lipid mobilization and oxidation as well as mitochondrial respiration during HFHS feeding.

### Pork and soy protein intake disrupt the hepatic AMPK-ACC axis in HFHS-fed mice

The transcriptional signature of altered lipid handling and mitochondrial respiration led us to examine the impact of dietary protein source on the master regulator of lipid oxidation and energy homeostasis, adenosine monophosphate kinase (AMPK), and its target acetyl-CoA carboxylase (ACC). Liver expresses two ACC isoforms: ACC1 is cytosolic and generates malonyl-CoA, classically viewed as destined for *de novo* lipogenesis, whereas ACC2 is associated with the outer mitochondrial membrane and produces malonyl-CoA in close proximity to carnitine palmitoyltransferase (CPT1), thereby restraining mitochondrial fatty acid import for beta-oxidation^24^. Pork and soy protein feeding resulted in a striking decrease in the ACC2 protein content in liver of mice fed the HFHS diet (Figure 2I, J). In contrast, total ACC protein levels were not significantly reduced by soy and pork proteins (Figure 2I, K), suggesting a preferential regulation of the ACC2 isoform impacting beta-oxidation flux. The decreased ACC2 protein content was not associated with changes in ACC phosphorylation at Ser79 (Figure 2I, L), despite increased AMPK activation based on T172 phosphorylation in pork protein-fed HFHS mice (Figure 2I, M). Given that ACC2-derived malonyl-CoA serves as a key allosteric inhibitor of mitochondrial fatty acyl-CoA import via CPT-1, these data suggest that soy and pork protein relieve this inhibitory break on mitochondrial fatty acid import, thereby promoting lipid flux into mitochondria independently of canonical ACC phosphorylation.

To distinguish whether this regulatory effect on ACC2 protein arises from a chronic adaptation to HFHS-induced obesity or from an acute, nutrient-driven response to dietary protein, we next conducted a fasting–refeeding experiment after a shorter, 2-week feeding period using the same diets as in the chronic protocol. Mice were then fasted for 12h and subjected to a 15-minute meal challenge on their respective diets, followed by tissue collection 2h post-refeeding (Figure 3A). Despite lack of body weight differences groups (Figure 3B), an even more robust reduction of ACC2 protein (and total ACC protein) for both soy and pork proteins fed mice on the HFHS diet was observed in this acute fasting-refeeding protocol (Figure 3C-E). Thus, downregulation of liver ACC2 protein is an early response to dietary protein source, rather than a secondary consequence to long-term exacerbation of obesity and hepatic insulin sensitivity. Consistent with the chronic study, pork protein feeding of HFHS mice for 2 weeks followed by refeeding increased AMPK phosphorylation (Figure 3C and F). Consequently, the higher ACC phosphorylation/ACC ratio in mice fed the soy and pork protein diets (Figure 3C and G) likely reflects the robust reduction in total ACC protein levels rather than increased kinase activity *per se*.

**Figure 3:**
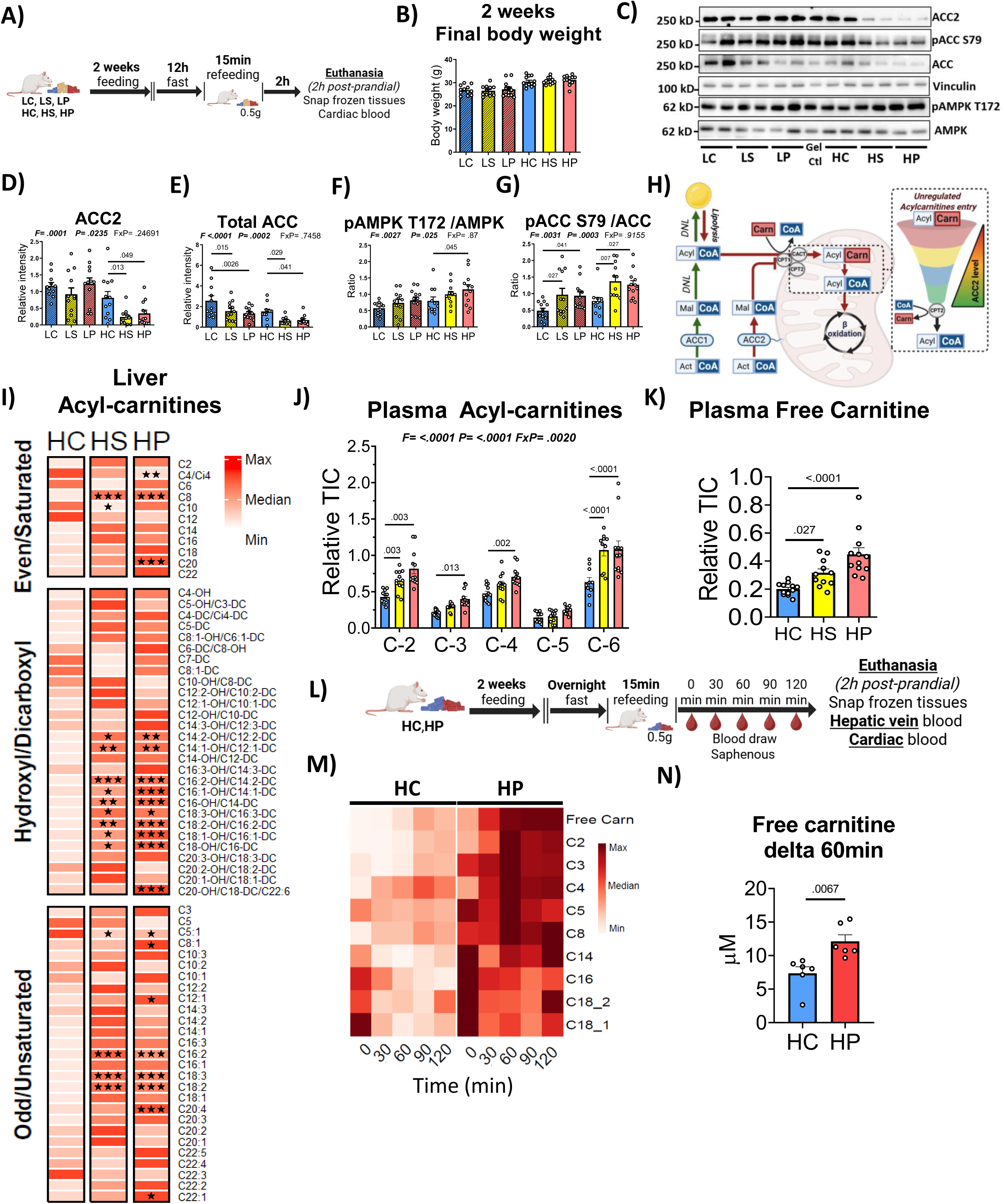
Pork and soy meal in HFHS induces postprandial mitochondrial lipid supply overload through ACC2 downregulation and increased carnitine delivery. **A)** Mice (n=12) were fed the 3 protein types in either low fat or high fat diet for 2 weeks, then were fasted overnight before a meal test of 15 minutes with 0,5g of diets to imitate a bolus. Mice were sacrifice 2h following the meal test, and blodd draw was taken every 30 minutes. **B)** Final body weight after 2 weeks of feeding. **C)** Representative gel for 2 mice per groups of ACC isoforms and AMPK phosphorylation. Gel Ctl= Gel loading control. **D-G)** Quantification of band intensity for proteins in B) for all mice (n=10-12) normalised on loading control vinculin and gel control. **H**) Schematic representation of the relationship between ACC2, the carnitine shuttle, mitochondrial beta-oxidation and acyl-carnitines. **I)** Liver acylcarnitine profile of mice (n=6) fed three dietary protein in high fat high sucrose liver 2h after a meal. **J-K)** Plasma level after 2h(n=6) of short (C-2 to C-6) chain acylcarnitine (E) and total free carnitine (F) in HFHS diet groups. **L)** Mice (n=12) were fed pork and casein protein in HFHS diet for 2 weeks, then were fasted overnight before a meal test of 15 min with 0,5g of diets to imitate a bolus. Blood draw was taken every 30 min from saphenous vein until sacrifice 2h after the meal. **M**) Detailed kinetic of apparition of free carnitine and acylcarnitine following a meal test (n=6 mice/group). **N)** Free carnitine delta from baseline (blood draw right after meal) to 60min after meal. Statistical analyses were performed using a two-way ANOVA, followed by a Dunnet’s post-hoc test for comparison to casein group. P-values of general effect for diet fat and sucrose (F), protein (P) are recorded under the title of each graph, followed by the p-values of the corresponding factor interaction effects.. Two tail Student t-test was used for panel N. Data are represented as means ± s.e.m, or color variation around the z-score applied to each metabolite for the heatmaps in M).

### Short-term consumption of pork and soy protein promotes post-prandial hepatic accumulation of long-chain acylcarnitines

Reduction in ACC2 is expected to diminish malonyl-CoA and relieve inhibition of CPT1 (Figure 3H). To test this hypothesis, we next measured post-prandial hepatic AC levels as a marker of hepatic beta oxidative fluxin 2-week HFHS-fed mice consuming different protein sources. Consistent with reduced ACC2 protein, inclusion of pork or soy protein in the HFHS diet led to robust hepatic accumulation of even-chain AC in the form of hydroxylated (OH), dicarboxylic (DC), and unsaturated intermediates (Figure 3I). OH species, in the form of acyl-CoA, are generated during beta-oxidation, and their reconversion to AC occurs when there is a bottleneck in beta-oxidation. Accumulation of these species is indicative of lower NAD+ bioavailability and redox imbalance^25^. This may result from the loss of ACC2 mediated AC oversupply, surpassing beta-oxidation capacity and NAD+ renewal. This mitochondrial fatty acid overload is a metabolic feature closely associated with insulin resistance^20,26^. In line with this interpretation, post-prandial hepatic acyl-CoA profiling revealed accumulation of multiple unsaturated long-chain species (e.g 18:1, 18:2, C17:1, C20:5, C19:3) as well as the medium chain acyl-CoA C8:1 in HFHS mice fed pork or soy protein (Supplementary figure 4). Together with our data revealing lower expression of genes coding for their beta oxidation, FADS1 and FADS2 (Figure 2H), our data suggest that pork and soy protein promote a bottleneck in lipid processing in mitochondria, particularly for the mono- and poly-unsaturated fatty acids, by increasing entry via CPT1 and decreasing beta oxidation efficiency.

Because even chain AC generated during beta-oxidation can be exported into circulation, we next assessed AC in plasma 2h after the meal. We observed a similar increase in even short chain AC, particularly C2 and C6 in plasma of pork and soy protein fed HFHS mice (Figure 3J). Free plasma carnitine levels were robustly increased in HFHS mice consuming dietary pork protein and slightly but also significantly increased by soy feeding (Figure 3K).

This observation is particularly relevant for pork protein fed mice, as dietary L-carnitine is abundant in muscle-derived foods and constitutes a major source of circulating carnitine^27–29^. To determine whether dietary carnitine availability alters the plasma AC pool in the post-prandial period immediately following a meal, we next examined the time-course of carnitine and AC changes in plasma over a 2-hour period following feeding HFHS mice with pork protein as compared to casein protein as depicted in Figure 3L. Total plasma free carnitine was higher at baseline (Figure 3M) and rose more dramatically following the pork protein meal, which is consistent with constant dietary exposure to the higher carnitine content of pork protein. We observed a clear switch from long-chain to short-chain AC in circulation following the meal, an effect that was also amplified by pork protein feeding. Free carnitine and even chain AC peaked in plasma at 60 minutes following feeding HFHS mice with pork protein (Figure 3M and N). A similar peak was observed for C3 and C5 acylcarnitine, which are derived from branched-chain amino acid oxidation. These metabolites remained robustly elevated up to 2h post feeding. Taken together, these results indicate that during HFHS feeding, dietary protein composition influences post-prandial hepatic lipid flux through higher provision of carnitine and downregulation of ACC2, leading to mitochondrial lipid overload, as reflected by the accumulation of both hepatic and plasma AC.

### L-carnitine supplementation *in vitro* recapitulates the effects of dietary protein on the hepatic AMPK–ACC axis and insulin signaling during obesogenic diet

Transport of long-chain fatty acids into mitochondria via CPT-1 requires carnitine to convert long-chain fatty acyl-CoA to their corresponding long chain AC (Figure 3H). We surmised that the effects of soy and pork protein on hepatic ACC2 may be mediated by increased carnitine availability, explaining how dietary protein impact hepatic mitochondrial fatty acid flux. To mechanistically address this hypothesis, Hepa 1-6 cells were exposed for 24h to either control medium (BSA) or a lipid- and sugar-rich medium containing BSA-conjugated palmitate, oleate and fructose (PA:OA:Fr), in the presence of increasing concentrations of L-carnitine (Figure 4A). Remarkably, carnitine supplementation induced a dose-dependent reduction in ACC2 protein levels, especially in the presence of the PA:OA:Fr mixture, reminiscent of the HFHS diet specific impact on ACC2 during the *in vivo* post-prandial protocol (Figure 4B and C). In line with the isoform specific regulation of carnitine on ACC2, the effect was less striking for total ACC protein (Figure 4B and D). We also observed a modest but significant effect of carnitine to increase AMPK phosphorylation in the PA:OA:Fr media but this effect was not present in cells cultured with BSA alone (Figure 4B and E). This is consistent with the *in vivo* effect of pork protein feeding; whereby liver AMPK phosphorylation was only observed in the context of HFHS feeding. Importantly, cells exposed for 24h to L-carnitine in the presence of the PA:OA:Fr media were also insulin-resistant as shown by the marked reduction of Akt Ser473 phosphorylation after acute insulin stimulation (Figure 4F and G). However, carnitine alone (in BSA condition) had no impact on insulin induced Akt Ser473 phosphorylation. These data support our model whereby increased post-prandial circulating level of carnitine coming from dietary protein exacerbates hepatic insulin resistance in the setting of HFHS feeding by removing ACC2 mediated regulation of mitochondrial beta oxidation, thereby promoting mitochondrial lipid overload.

**Figure 4:**
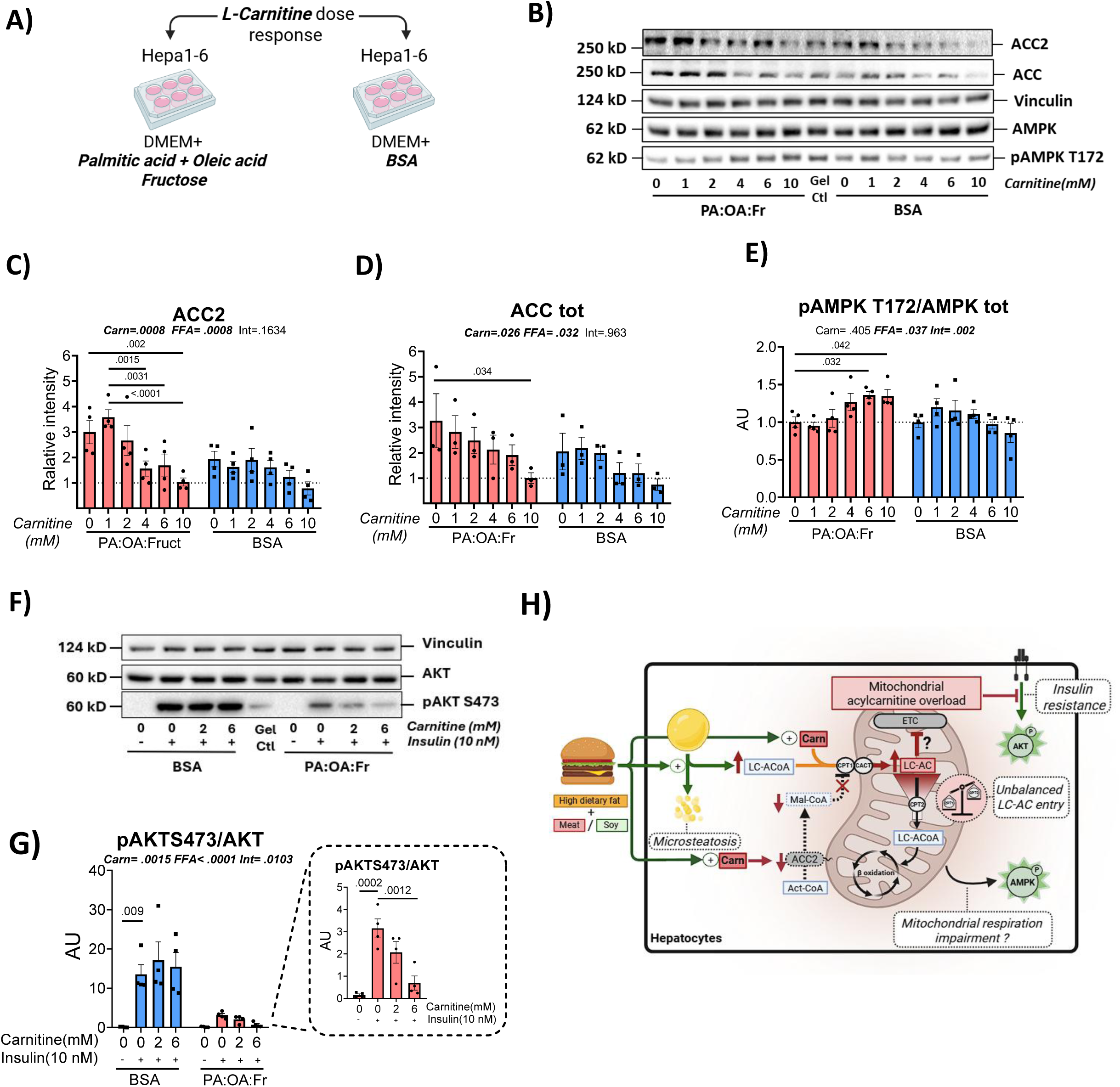
L-carnitine supplementation *in vitro*, combined with elevated long-chain free fatty acid levels, recapitulates the effects of dietary protein on the hepatic AMPK–ACC axis and insulin signaling. **A**) Hepa1-6 cells were treated for 24h with increasing dose of L-carnitine in high free fatty acid and fructose media (PA:OA:Fr = palmitate, oleate, fructose) or bovine serum albumin(BSA) supplemented media as control. **B)** Representative gel for the immunoblotting of the AMPK-ACC axis. Gel Ctl= Gel loading control. **C-D)** Quantification of level of total ACC2(C) and total ACC (D) protein in B) normalised on loading control vinculin and gel control. **E)** Quantification of level of activatory phosphorylation of AMPK on threonine172 normalised to total AMPK protein level as shown in B). **F-G)** Insulin sensitivity as shown by level of phosphorylated AKT after 10 minutes of insulin 10nM in 3h BSA deprived Hepa1-6. Cell were incubated for 24h with the indicated dose of carnitine in bovine serum albumin(BSA), fructose and free fatty acid rich media (PA:OA:Fr). Representative gel are shown in (F). **H)** Conceptual model of of proposed mechanism explaining how dietary protein interact with obesogenic diet to induce mitochondrial lipid overload and insulin resistance. Both the total amount of fat load and carnitine participate to the saturation of mitochondrial lipid oxidative capacity, which leads to AMPK activation as a stress signal to rebalance mitochondrial oxidative capacity. Data are represented as means ± s.e.m for 3-4 independent experiments done in technical triplicate. Statistical analyses were performed using a two-way ANOVA, followed by a followed by a Dunnet’s post-hoc test for comparison to the relevant control groups. P-values of general effect for carnitine (Carn), fatty acid and fructose (FFA) are recorded under the title of each graph, followed by the p-values of the corresponding factor interaction effects. Two-way ANOVA was done for insulin treated groups in F) and G).

## Discussion

The influence of dietary protein composition on metabolic dysfunction associated with obesity and T2D has garnered increasing attention in recent years^14,18,19,30–40^. We previously demonstrated that, compared to casein, a mixed dietary protein source impairs glucose homeostasis, increases post-prandial hepatic short-chain AC, and promotes insulin resistance, in part through alterations in gut microbiota^19^. However, gut microbiota composition does not explain the change in liver oxidative metabolism leading to the build-up of AC.

Here we identify a previously unrecognized mechanism linking dietary protein source to liver metabolism and insulin sensitivity. We show that elevated circulating carnitine, derived from specific meat protein sources such as pork, leads to a reduction in hepatic ACC2 protein, resulting in increased mitochondrial fatty acid influx, lipid supply overload, accumulation of AC, exacerbating systemic and hepatic insulin resistance in the context of HFHS feeding. This mechanism suggests that carnitine is not just a passive co-factor of the carnitine shuttle but actively regulates lipid oxidation through modulation of ACC2 abundance. This novel carnitine-ACC2 regulatory axis is distinct from classic kinase-based regulation of ACC, yet the precise molecular mechanism underlying the carnitine-dependent reduction of ACC2 remains to be fully elucidated. One potential explanation is that chronic activation of AMPK in HFHS-fed mice consuming pork protein led to ACC2 degradation as previously reported in mice constitutively expressing an active α-subunit, which led to a sharp decrease in total ACC protein in liver^41^.

The impact of pork protein on liver metabolism is reminiscent in many ways to the effect of hepatic double ACC1/ACC2 genetic ablation. Deja et al.^42^ reported reduced hepatic Akt phosphorylation in fast-refed mice lacking both liver ACC isoforms. Similar to us, they reported increased post-prandial short and medium chain AC in the liver of mice lacking hepatic ACC, even in chow-fed animals. Furthermore, liver of double ACC knock-out mice has increased redox imbalance and oxidative stress, again similar to the transcriptomic signature of oxidative stress responses we observed in liver of mice fed pork protein. Liver specific double ACC knock-out was also reported in another study to increase anti-oxidant defense and the levels of NADPH and reduced glutathione^43^. Likewise, we observed a significant increase in total glutathione in liver of mice fed pork protein (Supplementary Figure 3C).

On the other hand, it has been reported that genetic disruption^19,42–45^ or pharmacological inhibition^46,47^ of ACC reduce hepatic steatosis. We have not observed change in total liver steatosis and hepatic triglyceride levels in our HFHS-fed animals consuming different protein types. However, we found that both pork and soy protein intake reduces ACC2 protein in HFHS-fed mice in association with extensive microvesicular steatosis as compared to mice fed casein. The lack of changes in total steatosis in HFHS mice fed different protein types as compared to genetic or pharmacological models of ACC invalidation may be related to the fact that only ACC2 and not ACC1 was specifically impacted by dietary protein sources. Indeed, selective ACC2 genetic disruption in mice have a milder phenotype than liver-specific ACC1 deletion^48^. The ACC2 isoform is mainly involved in CPT1 inhibition and lipid oxidation, but less related to *de novo* lipogenesis^49^, which may explain our finding of specific modulation of microvesicular steatosis phenotype rather than global triglyceride storage.

Macrovesicular steatosis is a more common form of lipid accumulation in hepatocytes, while microvesicular steatosis is linked with a more advanced stage of steatosis, worsen liver function^50^ and steatosis grade^51^. Importantly, microvesicular steatosis is more closely linked with impaired mitochondrial β-oxidation, and an increase risk of progression to MASH^52^. Extensive microvesicular steatosis in pork protein fed animals might also be related to our finding of increased lipase activity and enhanced access to the surface of the lipid droplet. We observed more NEFA accumulation in HFHS mice fed pork and soy proteins as compared to casein, suggesting a disrupted balance between lipid mobilization and mitochondria oxidative capacity caused by lipid excess in the former groups. It is interesting to note that a recently proposed role of lipid droplets is to serve as a sink to prevent harmful oxidation of phospholipids, especially PUFA that are sensitive to oxidative stress^53^. As many PUFAs accumulated in the form of acyl-CoA or AC in the liver of pork and soy protein fed mice during HFHS diet, it is possible that microvesicular steatosis is a consequence of impaired cellular response to PUFA peroxidation, which could ultimately impact insulin sensitivity and liver glucose metabolism.

An emerging feature of dietary protein’s impact on energy metabolism and insulin action is its modulatory effect on plasma and liver AC, as we previously reported^19^. We now propose that this mechanism at least partly involves the provision of carnitine from specific dietary protein sources, such as pork protein, that lowers liver ACC2 content and releases malonyl-CoA inhibition of the CPT1a shuttle, leading to unregulated entry of FFA-CoA into the mitochondria for β-oxidation. This is supported by the levels of liver AC, coupled with transcriptomic data, which is consistent with an impaired mitochondrial oxidation capacity in the face of increased lipid influx in the context of HFHS feeding. In contrast to meat, plant-based protein sources such as soy do not contain carnitine yet we did find elevated systemic carnitine levels in soy protein HFHS fed mice. This may result from *de novo* biosynthesis of carnitine, which is the predominant mechanism of carnitine bioavailability in herbivores but also in strict vegetarians and vegans^54,55^, and which has been recently reported to be mediated by a novel mitochondrial carrier of trimethyllysine (SLC25a45)^56^.

There is growing evidence that mitochondrial lipid overload linked with high beta-oxidation rates can lead to insulin resistance^26,57^. The direct effect of ACs, particularly the long-chain species, on insulin resistance and mitochondrial respiration has been reported in muscle and fat cells *in vitro*^58–60^ and mouse skeletal muscle^61^, but not in liver. We found that carnitine supplementation of Hepa 1-6 cells incubated with high FFA-Fructose load also promoted insulin resistance, as revealed by impaired Akt activation by acute insulin, similar to what was previously observed in myocytes^26^. In heart, AC can increase electron leakage and ROS production^62^, a recognized mechanism of insulin resistance. Our transcriptomic results indicate that dietary pork protein, and to a lesser extend soy protein, activates oxidative stress buffering genes in HFHS diet, which aligns with increased ROS production. It is however unclear if accumulation of ACs is merely a reflection of high levels of lipolytic intermediates, like diacylglycerol and ceramides, which could account for impaired insulin signaling^63^.

Long-chain fatty acyl-CoA can directly activate AMPK in hepatocytes through allosteric modification^64^. Our results show that AMPK activation in HFHS-fed mice occurs only when mice were fed dietary pork protein which had higher levels or plasma and hepatic carnitine. This effect was mimicked through direct media supplementation of carnitine to Hepa 1-6 cells incubated with a mix of fatty acids and fructose. It should be noted that supplementing L-carnitine to hepatic cells without the presence of fatty acids and fructose (BSA condition) did not increase AMPK phosphorylation, indicating that L-carnitine is not a direct activator of AMPK in liver cells. This suggests that carnitine activates AMPK only during mitochondrial lipid overload, a condition linked with respiration inefficiency and elevated reactive oxygen species (ROS) level. This is consistent with the fact that in human fibroblasts, long-chain AC dose-dependently reduce OXPHOS-dependent mitochondrial respiration, leading to increased generation of ROS and reduce ATP to AMP ratio^65^. Accumulation of ROS has been shown to directly increase AMPK activity through oxidation of cysteine residues, inducing conformational changes that enhance its susceptibility to phosphorylation at threonine 172^66,67^.

This study has important translational implications for human nutrition and health. While red meat consumption is linked to metabolic and liver disease incidence^68–72^, leading to recent European health recommendations to reduce red meat intake^73^, the molecular mechanisms underlying this relationship remains elusive. Saturated fats in red meat are often blamed^68,74,75^. We have, however, controlled for total lipids and saturated fat levels in our experimental diets, and still observed major changes in systemic and liver metabolic impairments. These observations point towards a key and independent role for dietary protein composition in regulating glucose homeostasis and liver metabolism, in the context of excess fat and sugar intake. Our pre-clinical observations may be particularly relevant for guiding nutritional recommendations about the type of dietary protein to consume, rather that only focusing on how much protein we should eat, for preventing obesity and better management of T2D and MASLD risk. Our results clearly show that, at least in animal models of diet-induced obesity, the type of protein sources consumed is important and that animal (meat) and even plant-based protein sources can exacerbate weight gain and T2D. In this regard, it appears that dairy proteins, as represented by casein here, represent a better protein source owing to the lack of dietary carnitine provision or *de novo* carnitine biosynthesis. This could contribute to the well documented neutral or even preventive effects of dairy products on obesity, T2D and MASLD in large-scale epidemiological studies^76,77^. Similar to animal protein, this negative relationship with T2D incidence is generally attributed to the lipids contained in dairy foods (e.g. C14:0 sphingomyelin and C34:0 phosphatidylethanolamine). The low carnitine level in this protein source should be given more attention, especially since it is also associated with lower circulating levels of γ-butyrobetaine^78^, a pro-atherogenic intermediate metabolite derived from L-carnitine^79^.

Our findings are also relevant to the current debate about using ketogenic diets to alleviate metabolic disease risks^80–85^. Indeed, recent clinical trials reported that low-calorie ketogenic diets can rapidly improve liver steatosis and insulin sensitivity^86,87^. However, the beneficial effects on liver steatosis are likely explained by substantial caloric restriction and rapid weight loss whereas the long-term impact of this diet on liver health remains unknown. This is of clinical importance since it was reported that key markers of liver injury such as the NADH/NAD+ ratio and TCA cycle turnover are changed in individuals fed a ketogenic diet, raising concerns on the long-term consequence on liver health and insulin sensitivity^86^. Accordingly, preclinical studies controlling for caloric intake reported liver-specific inflammation and insulin resistance despite weight loss with ketogenic diets^88–90^. One may speculate that simply changing the protein source could mitigate some the long term hepatic adverse effect of such diets^91^.

In summary, we identified a novel mechanism by which dietary protein source influences obesity, glucose homeostasis and hepatic metabolism. We propose that dietary carnitine drives a previously unrecognized regulatory axis involving ACC2, leading to mitochondrial lipid oversupply surpassing oxidative capacity, resulting in exacerbating insulin resistance during HFHS feeding. These results highlight the importance of considering how protein source, rather than quantity, interacts with dietary carbohydrate and lipid intake to alter liver metabolism, and ultimately influence obesity and T2D progression.

## Method

### Diets

The following diets were elaborated: low-fat low sucrose (LFLS: 70% kcal carbohydrates, 7% kcal sucrose, 15% kcal fat) and high-fat high-sucrose (HFHS: 35% kcal carbohydrates, 28% kcal sucrose, 50% kcal fat) diets containing 15% kcal of dietary protein comprised of either powdered lyophilized pork (Happy Yak, Canada), isolated soy protein (Teklad Envigo, USA) or isolated casein (MP Biomedicals, USA). Cuts of meat came from different parts of the animal without any additional processing beyond lyophilization. The composition of each protein source has been analyzed (Environex) and was used to normalize macro and micronutrients in the different diets (Supplementary Tables 1 and 2). Differences in fat amount in different protein sources were compensated with addition of lard fat or corn oil to match a ratio of saturated to polyunsaturated fatty acids (SAT:PUFA) of 1.33, representing the daily American dietary intake^92^. Fiber content was matched to 5% (w/w) with alpha cellulose. For carbohydrate calculation, corn starch (MP Biomedical, USA) was adjusted based on 91% amidon, 9% humidity. Sucrose addition was also adjusted for the Vitamin mix (AIN-76A, Tecklad, USA) containing 98% (w/w) sucrose, and Mineral mixture 76 (MP biomedical, USA) containing 12% sucrose. Final composition in %kcal is summarized in Supplementary Table 3.

### Animal protocols

All protocols were carried out with C57BL/6J male mice single-housed in ventilated cages at a humidity of 40–50% and a temperature of 22 °C on a 12-hour dark-cycle with ad libitum food. 6-week-old mice were used for the 12-week protocol whereas 10-week-old mice for the 2-week protocol. Mice were acclimatized for 2 weeks on the LFLS-C diet prior to the beginning of their respective treatment. Animals were then distributed into treatment groups according to their body weight and fed with their respective diets for 12 weeks or 2 weeks. Food intake was measured three times a week and body weight weekly. Body composition was measured by nuclear magnetic resonance with a Bruker Minispec (LF90) apparatus.

For the 12-week protocol, a fasting blood sample was collected at baseline and at week eleven. A blood sample in the fed state was collected at week 6. Intraperitoneal insulin tolerance test (ipITT) was performed at week 10 with 0.65 U/kg insulin dose injection after a 6h fast, and tail vein sampling after 10, 20, 30 and 60 min. Oral glucose tolerance test (oGTT) was performed at week 12 and after a 6h fast using a dose of 1mg of glucose/g of BW, with tail vein sampling after 15, 30, 60 and 120 min. Animals were euthanized at week 12 after 6h fast, and just following an acute (5 min) intravenous injection of saline or insulin (3.8 U/kg) for later analysis of hepatic insulin signaling. Portal vein was sampled first, followed by cardiac puncture. Liver was rapidly weighted, the frontal lobe was snap frozen using pincers pre-cooled in liquid nitrogen, while the other lobes were rapidly collected for histology (OCT and PFA). Other tissues (eWAT, BAT, rpWAT, pancreas, mWAT, gastrocnemius muscles) were weighted and snap frozen in liquid nitrogen.

For the 2-week protocol, a meal test was conducted at the end of the dietary treatments, where 0.5 g of their respective diet was given to the animals after a 12-h fast. Mice had 15 min to eat the food and were then euthanized 2 h after the end of the meal test. Portal vein was sampled first, then liver front lobe was quickly snap frozen as above.

All animals were euthanized after portal vein sampling by cardiac puncture (under isoflurane anesthesia) and cervical dislocation before tissue collection for analysis. All manipulations were approved (VRR #20-421 and #21-842) by the Comité de protection des animaux de l’Université Laval (CPAUL) and complied with the Canadian Council on Animal Care (CCAC) guidelines.

### Biochemical analyses

Blood glucose was recorded using a One Touch glucometer UltraMini® and values higher than the upper limit of the glucometer were determined by an Amplex® red glucose/glucose oxidase assay kit (Sigma-Aldrich, USA). ELISA assays were used to measure insulin (Alpco Mouse Ultrasensitive Insulin kit). Cytokines and diabetes related metabolites were measured in liver lysate using Bio-Rad Multiplex Immunoassays with Luminex xMAP^®^ Technology separating IL-1b, IL-10, KC, TNFa, MCP1 and Resistin and PAI-1 in two different panels. Lipase activity was measured with a fluorometric assay using the Lipase Detection Kit III (#ab118969, Abcam, USA) following the manufacturer’s instructions. Liver triglycerides, cholesterol, and NEFA were extracted by an adapted Folch method^93^ and dosed with colorimetric kits (Infinity^TM^ for triglycerides, WAKO HR series NEFA-HR for NEFA and Randox Life Science for cholesterol). Reduced and oxidized glutathione was measured using Glutathione Assay Kit (Fluorometric) (ab65322, Abcam, USA).

### Acyl-CoA and Acylcarnitine LC-MS analysis

For Acyl-CoA quantification, 50 mg tissue samples were homogenized in 1ml extraction buffer (1:1 methanol/water containing 5% acetic acid) followed by a 15-minute centrifugation at 18,000g. For solid phase extraction, the cartridge (1 ml ion exchange cartridge packed with 100 mg of 2-2(pyridyl)ethyl silica gel (Sigma)) was pre-activated with 1 ml of methanol and then with 1 ml of extraction buffer (1:1 methanol/water containing 5% acetic acid). The acyl-CoAs trapped on the silica gel cartridge were released with (i) 1 ml of a 1:1 mixture of 50 mm ammonium formate, pH 6.3, and methanol and then (ii) 1 ml of a 1:3 mixture of 50 mm ammonium formate, pH 6.3 and methanol and finally (iii) 1 ml of methanol. The combined effluent was dried with nitrogen gas and stored at 80°C until LC-MS analysis^57^.

For Acyl-Carnitine, 20 mg frozen and pulverized tissue was spiked with 20 ml 0.01 mM D9 carnitine as internal standard and homogenized with a Folch-extraction approach. A 300 ml tissue extract was completely dried by nitrogen gas. The derivatized sample was dried again by nitrogen gas and the dried sample was re-dissolved into 20 ml methanol followed by 60 ml LC water. The derivatized samples were analyzed by LC-QTRAP 6500+-MS/MS (Sciex, Concord, Ontario). A 5 ml sample was injected on a Pursuit XRs 5 C18 column (150 x 2.0 mm, 5um), protected by a guard column (Pursuit XRs 5 C18, 10x 2.0 mm, 5μm) with temperature controlled at 25°C, in an ExionLC AD liquid chromatograph system. The chromatogram was developed at 0.4 ml/min (i) for 2 min with 100% buffer A (98% H2O and 2% acetonitrile with 0.1% formic acid), (ii) from 2 to 13 min with a gradient from 0 to 80% buffer B (98% acetonitrile and 2% H2O with 0.1% formic acid), (iii) from 13 to 18.5 min with 10% buffer A and 90% buffer B, (iv) from 18.5 to 19 min with a gradient from 90% to 0 buffer B, and (v) for 10 min of stabilization with 100% buffer A before the next injection. The liquid chromatograph was coupled to a 6500+-MS/MS operated under positive ionization mode with the following source settings: turbo-ion-spray source at 650°C under N_2_ nebulization at 65 PSI, N_2_ heater gas at 65 PSI, curtain gas at 35 PSI, collision-activated dissociation gas pressure held at high, turbo ion-spray voltage at 5,500 V, declustering potential at 33 V, entrance potential at 10 V, collision energy ranged from 24 to 46 V optimized for each acylcarnitine and collision cell exit potential at 14 V. The Analyst software (version 1.6) was used for data recording and processing.

For plasma Acyl-Carnitine and free carnitine, 40μl of plasma was homogenized in 150 µl of frozen acetonitrile (−20 °C) containing 0.2% formic acid and spiked with a mixture of labeled internal standards: 1.25 nmol of d9-Car, 250 pmol of d3-C2 Car, 10 pmol of d3-C3 Car, 5 pmol of d3-C4 Car, and 2.5 pmol of d3-C8 Car. A zirconium oxide ceramic bead (2.8 mm; VWR, Mississauga, ON, Canada) was added, and the sample was homogenized using a Bioprep-24 Homogenizer (Montreal Biotech Inc., Montreal, QC, Canada) for 20 seconds at low intensity. The mixture was then vortexed for 2 minutes and centrifuged for 10 minutes at 12,000 × g. The resulting supernatant was transferred to an LC vial for analysis. Samples were analyzed using a 1290 Infinity HPLC system coupled to a 6545 accurate-mass quadrupole time-of-flight mass spectrometer (LC-QToF; Agilent Technologies) equipped with a Dual Agilent Jet Stream ESI source. Separation was performed using a PEEK-coated SeQuant ZIC-pHilic guard column (20 × 2.1 mm; Millipore, Billerica, MA, USA) and an Atlantis HILIC Silica column (3 µm, 2.1 × 150 mm; Waters, Milford, MA, USA). The mobile phase consisted of solvent A (10 mM ammonium formate with 0.1% formic acid) and solvent B (acetonitrile with 0.1% formic acid). Samples were eluted over 26 minutes at a constant temperature of 35 °C and a flow rate of 300 µl/min using the following gradient: 0–2 min, 90% B; 2–15 min, 90–50% B; 15–16 min, 50–10% B; 16–19 min, 10% B; 19–20 min, 10–90% B; followed by a 6-minute re-equilibration. LC-MS operating parameters were as follows: gas source temperature, 250°C; drying gas flow rate, 10 L/min; sheath gas temperature, 250°C; sheath gas flow rate, 8 L/min; nebulizer pressure, 35 psi; capillary voltage, 3500 V; and nozzle voltage, 1500 V. Continuous mass correction was applied during LC-MS analysis via on-line infusion of purine. Mass spectra were acquired in positive electrospray ionization mode over an m/z range of 20–300, with MS scans collected every 1 s.

### Hepatic histology

Formalin-fixed paraffin embedded liver samples were cut (4 μm) and stained with hematoxylin and eosin (H&E) on an automatized platform for histologic assessment. Liver sections were blindly evaluated by 3 individuals and revised by a pathologist at the IUCPQ research center using the NAFLD scoring system proposed by Liang et al^94^. The percentage of hepatocytes harboring either macro- or microvesicular steatosis, or hypertrophy (increase of more than 1.5x the size of a normal hepatocyte) was scored as steatosis. Inflammation was evaluated as the number of inflammatory foci (cluster of at least five inflammatory cells) per field at 10x magnification (3.5 mm^2^) in at least five fields. The presence of steatosis in more than 5% of hepatocytes was classified as NAFLD and the association of both steatosis and inflammation in more than 0.5 cluster per field was classified as NASH. Other paraffin-embedded sections (6 μm thick) were deparaffinized through toluene and a graded alcohol series to assess liver fibrosis. Sections were stained for 8 min with Weiger Hamatoxyline and rinsed for 5 min in water. Subsequently, the sections were stained for 60 min with Sirius red (0.1% of Sirius red in saturated aqueous picric acid) for collagen bundle staining. Sections were then rinsed with acetic acid 0.05% and mounted for observation under polarized light microscopy (Zeiss AXIOPLAN; Oberkochen, Germany). Fibrosis was scored according to the definition used by Kleiner et al. A score of 1 was given when fibrosis was found in perisinusoidal or periportal regions, 2 when fibrosis was present in both, 3 with bridging fibrosis, and 4 with cirrhosis.

### Bulk RNAseq

Library preparation was performed by Genome Québec (Montreal,Canada) with 10 mg of total RNA with a Bioanalyzer RIN score greater than 8.0. Ribosomal RNA was removed by poly-A selection using Oligo-dT beads (mRNA Direct kit, Life Technologies). mRNA was then fragmented in a buffer containing 40 mM Tris Acetate pH 8.2, 100 mM Potassium Acetate, and 30 mM Magnesium Acetate and heated to 94 degrees for 150 seconds. mRNA was reverse transcribed to yield cDNA using SuperScript III RT enzyme (Life Technologies, per manufacturer’s instructions) and random hexamers. A second strand reaction was performed to yield ds-cDNA. cDNA was blunt ended, had an A base added to the 3’ ends, and then had Illumina sequencing adapters ligated to the ends. Ligated fragments were then amplified for 12 cycles using primers incorporating unique index tags. Fragments were sequenced on an Illumina HiSeq-3000 using double reads extending 50 bases.

Both reverse and forward stranded reads were done with mean read lenght of 101 base pairs. Cleaning of the fastQ files, alignment and counts were done on Galaxy server (https://usegalaxy.org/) using respectively Trimmomatic, RNA STAR and FeatureCounts on exon. Statistical analysis were done on Rstudio (V2026.01.2-418) using DESeq package from Bioconductor. ‘Protein’ (pork and casein) and ‘FatSucrose’ (HFHS or LFLS) were given as the two factor with two level using the design argument *design=∼FatSuccrose+Protein+FatSucrose:Protein.* Pathway analysis was done using Goseq package with org.Mm.eg.db mouse genome annotation version 3.22.0. Heatmaps were generated using “pheatmap” package and bubble plots were done using ggplot2 package. Significant genes (adjusted p-value <0.1) that had the occurrence “Fatty acid” or “Glycerolipid” in their KEGG annotation database were selected and we expressed the normalized gene count for all the dietary groups. These genes were ranked from the most upregulated at the top to the most downregulated at the bottom based on their log2 fold change. We also ran the same analysis comparing the subset of HFHS fed groups. The complete list of significant genes for each of these analyses can be found in Supplementary tables 4 to 7.

### Immunoblotting

Liver(30ug) were powdered with liquid nitrogen and then homogenized under rotation for 2h at 4°C in a 10-fold mass excess of ice-cold RIPA lysis buffer (50mM Hepes pH 7.5, 150mM NaCl, 1mM EGTA, 20mM β-glycerophosphate, 1% NP-40, 10mM NaF, 2mM Na3VO4, 0.1mM PMSF and protease inhibitor cocktail). Lysates were clarified by centrifugation at 16,000 × g for 10 min at 4°C and the proteins were measured with a BCA assay (ThermoFisher Scientific, Burlington, Canada). Cells were washed with PBS and lysed in ice-cold RIPA lysis buffer and were processed similarly to tissue lysis described above. Tissue and cell lysates were denatured in SDS sample buffer and submitted to SDS-PAGE followed by transfer on nitrocellulose membranes for immunoblotting (Pall Corporation, Mississauga, Canada). Membranes were blocked for 1h at room temperature and then probed with the primary antibody overnight at 4° C. After washing in TBST (50mM Tris-HCl pH 7.5, 0.15mM NaCl, and 0.1% Tween-20), the membranes were incubated with HRP-conjugated secondary antibodies for 1 h at room temperature. The detection was performed with an ECL reagent (Millipore, Etobicoke, Canada). Antibodies used were ACC2 (Mybiosource #MBS248196, 1/2000 in 2,5% BSA), ACC total (Cell signaling #3662, 1/10 000 in 2,5% BSA), phosphorylated ACC Serine 79 (Cell signaling #3661, 1/1000 in 2,5% BSA), Actin (Sigma #MAB150,1/50 000 in 2,5% BSA), Vinculin (Cell signaling #4650, 1/5000 in BSA 2,5%), phosphorylated AKT serine 473 (Cell signaling #9271, 1/1000 in BSA 5%), AKT total (Cell signaling #9272, 1/2000 in BSA 5%), phosphorylated AMPK Threonine 172 (Cell signaling #2335, 1/1000 in BSA 5%), AMPK total (Cell signaling #2532, 1/2000 in BSA 5%).

### Cell culture

Hepa 1-6 cells were thawed at passage 3-5. For carnitine supplementation experiments, cells were grown for 24h in DMEM +10% FBS, then cell media was changed for either a LFLS resembling media (BSA 100uM), or a HFHS resembling media, composed of high levels of free fatty acids (palmitic acid and oleic acid) and fructose (PA:OA:Fruct). The fatty acid mix of oleate to palmitate with a 1:1 ratio was mixed in ethanol and this FFAM was precomplexed to BSA in a ratio of 4:1 (4mM solution of FFAM to 1mM BSA, with an ethanol concentration of 1%). The final concentration of FFAM was 400 uM with 0.1% ethanol. Fructose’s final concentration was 15mM and carnitine was added to the concentration mentioned in the figures. Cells were incubated with these treatment media for 24h, then washed with ice-cold PBS and lysed for immunoblotting using RIPA lysis buffer as described above.

For insulin signaling experiments, after 21 hours of treatment, cells were incubated for 3h in media without FBS, but with the same treatments. After 3h of FBS deprivation, cells were stimulated with 10nM insulin for 10 minutes, then lysed for immunoblotting.

### Statistical analysis

Data are expressed as means ± s.e.m. Normality was assessed using Shapiro-Wilk’s test. For the 12-week and 2-week protocol, a two-way or a three-way analysis of variance (ANOVA) was performed to isolate the main effects of high fat diet (F), protein (P) or time for ITT and GTT (T), as well as the corresponding interaction effects. A Dunnet’s post-hoc test with a type I error set at 0.05 was then applied when the p < 0.10 threshold of main factors was achieved.

The exact p-values for main effects are indicated under the title of each graph and subsequent significant differences established by post-hoc test for specific group comparison are reported as p-value directly on graph. Differences in parameters involving repeated measures were assessed using a mixed linear model Rstudio, with treatment, time (T) and treatment*time interaction as fixed effects and an autoagressive covariance matrix to account for within-subject correlations. The skewness in the distribution of model residuals was considered, and data were log-transformed when required.

For metabolomics (high number of metabolites with low sample size) repeated one-way ANOVA was done for each metabolite and then corrected by Benjamini–Hochberg FDR to account for multiple testing. For RNAseq analysis, DESeq normalization and FDR were applied, and only adjusted p-values were used for selection of significant genes. Microsoft Excel (v16.47.1) and Graphpad Prism (v9.1.0) software were used to compile data and make graphs and figures.

## Data availability

All data generated or analyzed during this study are included in this article (and its supplementary information files, for complete RNAseq data). Metabolomics data will be uploaded to the right depository upon publication.

## Acknowledgement

We thank Joanie Dupont-Morissette Christine Dallaire, Bruno Marcotte for excellent technical assistance with animal protocols. We also thanks Noemie Daniel and Béatrice Choi for diets design assistance.

## Contributions

**Conceptualization:** F.B., A.M., **Methodology:** F.B, A.M., P.J.W.,S.D **Diet Design:** F.B, A.M, W.G **Investigation - In Vivo Analysis:** F.B., L.R.P, M.Sc, B.B., **Investigation – In Vitro Analysis:** F.B. **Data Integration:** F.B., P.M. **Writing - Original Draft:** F.B, **Writing - Review & Editing:** A.M., P.J.W, C.D.R, S.D., A.C. **Supervision:** A.M, A. C. **Approval of Final Manuscript:** All authors.

## Correspondence

and requests for materials should be addressed to André Marette.

## Legends to figures

**Supplementary figure 1:**
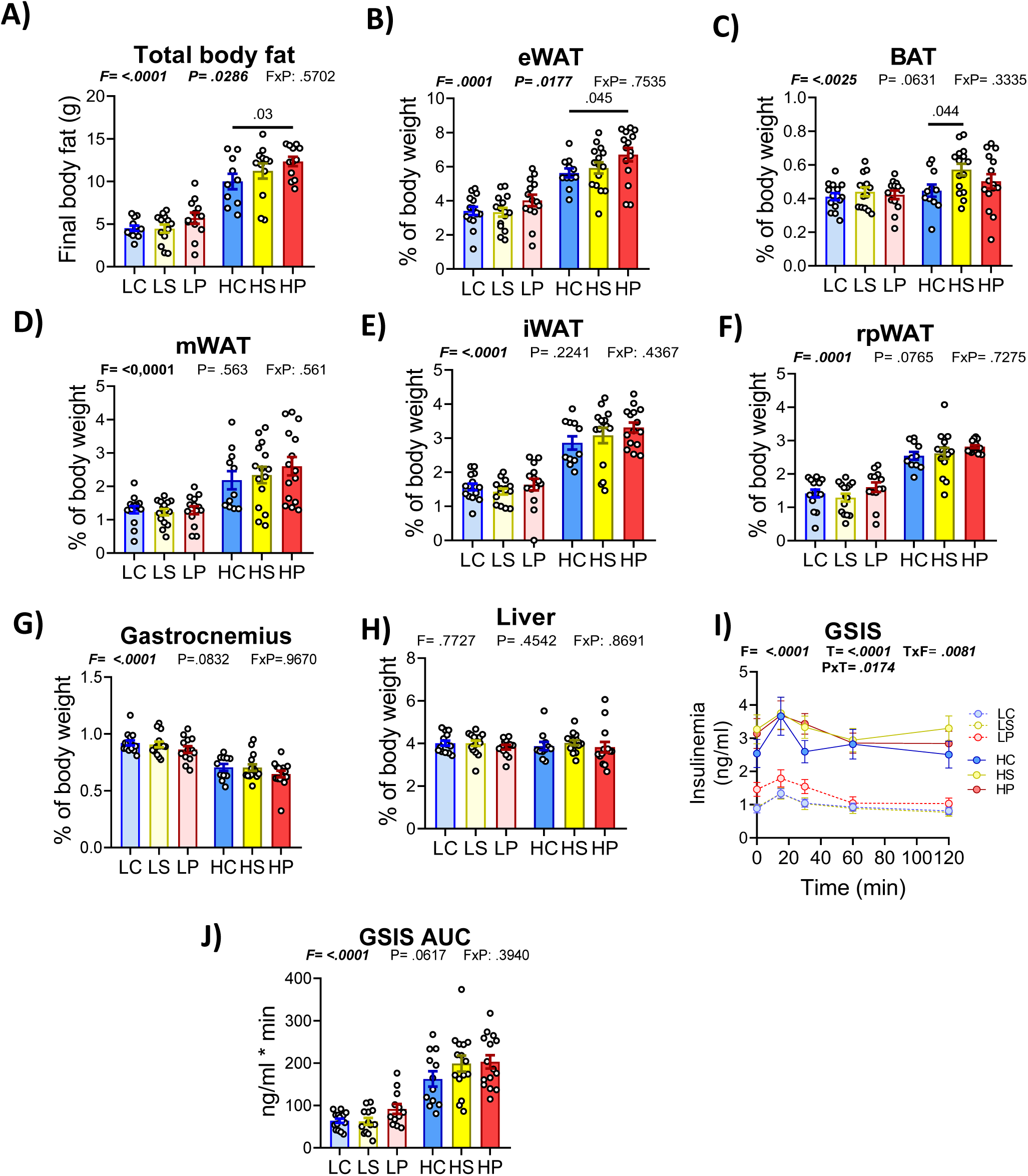
Impact of different dietary protein in either a low-fat low sucrose of high fat high sucrose on glucose stimulated insulin secretion and body composition after 12 weeks. **A)** Final body fat in grams as determined by TD-NMR **B**) Percentage of body weight from epididymal white adipose tissue **C-I)** Tissue weight was measure shortly after the sacrifice before being snap frozen in liquid nitrogen.BAT=Brown adipose tissue, mWAT=mesenteric white adipose tissue, iWAT= inguinal white adipose tissue, rpWAT= retro-peritoneal white adipose tissue. **J-K**) Glucose stimulated insulin secretion(GSIS) as well as respective area under the curve starting at baseline insulinemia (K) was measured by ELISA using tail vein blood collected during oral glucose tolerance test performed after 6h fast during the 12^th^ week of the protocol. Data are represented as means ± s.e.m from 12-15 mice per group. Statistical analyses were performed using a two-way ANOVA, followed by a Tukey post-hoc test. P-values of general effect for diet fat and sucrose (F), protein (P), time(T) are recorded under the title of each graph, followed by the p-values of the corresponding factor interaction effects. Detailed significant differences detected by post-hoc test are recorded as follows: *p < 0.05, **p < 0.01, ***p < 0.001.

**Supplementary figure 2:**
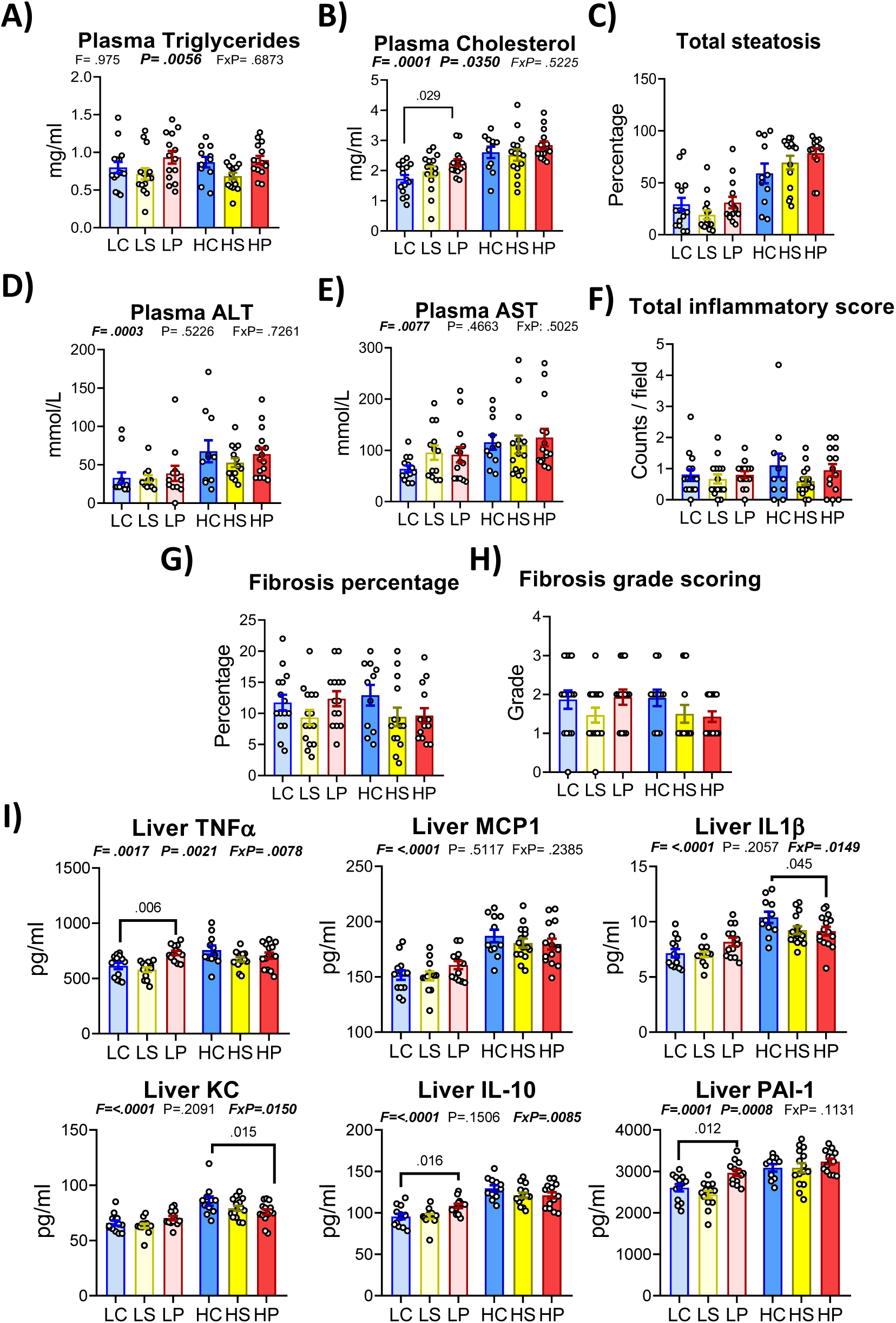
Impact of dietary protein source on circulating triglycerides and cholesterol and liver steatosis marker. **A-B)** Circulating triglycerides (A) and cholesterol B) assessed by colorimetric assay. **C)** Mean of total steatosis score from blinded visual quantification of three observers. **D-E)** Circulating liver transaminase AST and ALT in plasma from cardiac punction at sacrifice. **F)** Total inflammatory score based on the mean number of inflammatory foci per field for 3 different field per slide, with more than 3 immune cells cluster being counted as inflammatory foci. **G-H)** Sirius red staining was used to quantify fibrosis percentage(G). The scoring (H) was based on the amount and location along the perisinusoidal space form portal to central vein. **I)** Liver cytokines, diabetes and fibrosis marker measured by multiplex immunoassay. Data are represented as means ± s.e.m from 11-15 mice for the 12 weeks project. Statistical analyses were performed using a two-way ANOVA, followed by a Dunnet’s post-hoc test for comparison to casein group. P-values of general effect for diet fat and sucrose (F), protein (P) are recorded under the title of each graph, followed by the p-values of the corresponding factor interaction effects..

**Supplementary figure 3:**
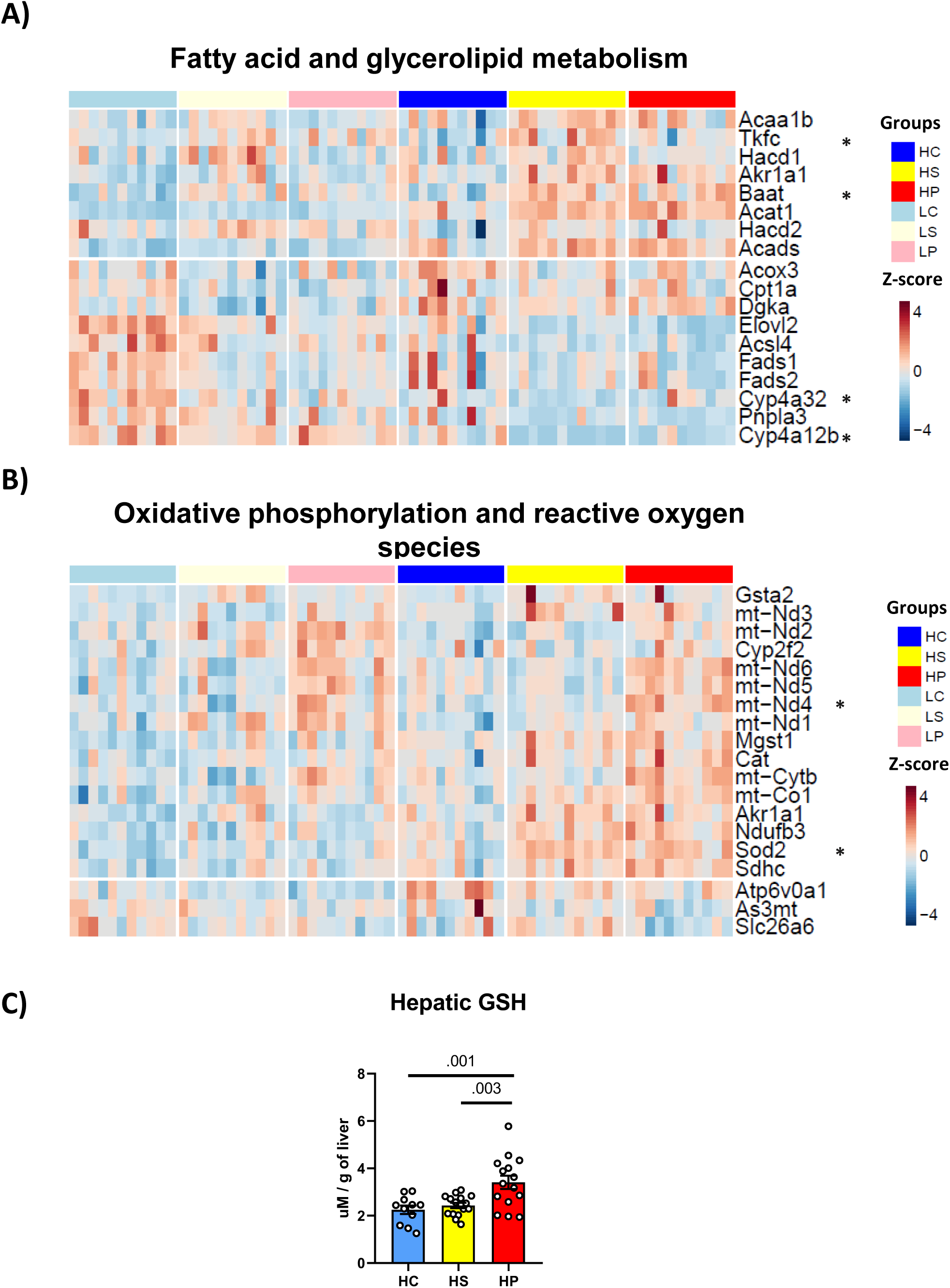
Pork and soy protein feeding induce transcriptional response related to mitochondrial respiration and fatty acid metabolism. **A)** Trimmed Mean of M-values (TMM, normalised feature counts) were used to visualise gene expression for significant differently expressed gene(padj<0.1) found during DESeq analysis for Soy to Casein comparison(overall protein effect, adjusted for diet background). Selection was made for gene with tag “Fatty acid” or “glycerolipid” in their KEGG annotation, and stars (*) indicates that these genes were significant for the subset HFHS-Soy to HFHS-casein comparison. **B)** Similar analysis as in A), this time comparing pork to casein and selecting genes based on the tags tag “oxidative phosphorylation” or “reactive oxygen species” in their KEGG annotation. Stars (*) indicates that these genes were significant for the subset HFHS-Pork to HFHS-casein comparison. **C)** Total glutathione in liver of HFHS groups after 12 weeks of feeding. One-way ANOVA with Dunnett’s multiple comparison test’s p-value are indicated from casein comparison.

**Supplementary figure 4:**
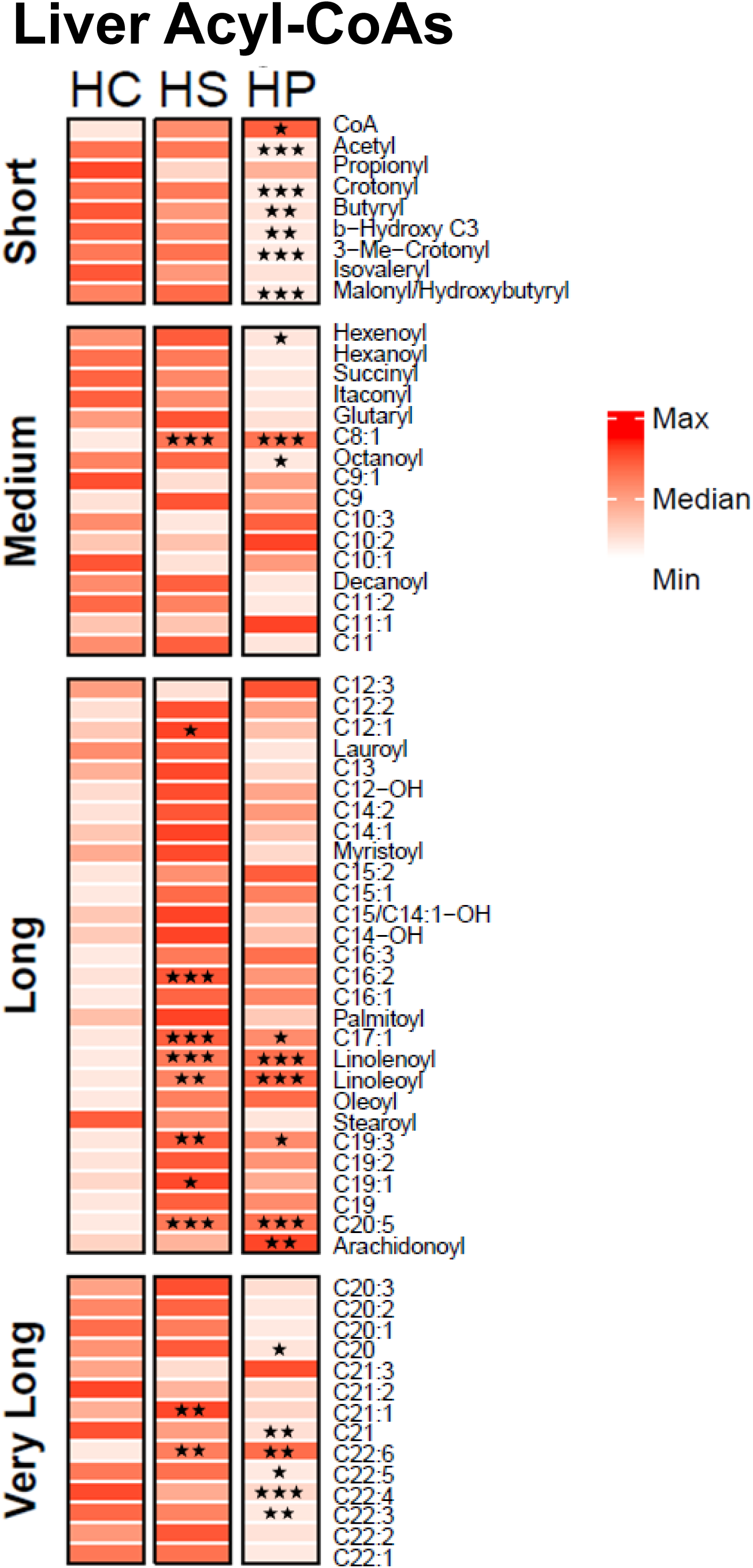
Liver acyl-Coa profile is altered after a meal of HFHS soy and pork protein. Acylcoa profile in liver of mice (n=6) fed three dietary protein in high fat high sucrose liver 2h after a meal. Each species were normalised using z-score transformation. Data are represented as color variation around the z-score applied to each metabolite for the. Statistical analyses were performed using repeated 1-way Anova for all metabolites, then corrected by Benjamini–Hochberg FDR to account for multiple testing. Detailed significant differences detected by post-hoc or t test are recorded as follows: *p < 0.05, **p < 0.01, ***p < 0.001.

